# Sequence Determinants of Human Junctophilin-2 Protein Nuclear Localization and Phase Separation

**DOI:** 10.1101/2021.05.17.444513

**Authors:** Ang Guo, Wenjuan Fang, Savannah Gibson

## Abstract

Junctophilin-2 (JPH2) was conventionally considered as a structural membrane binding protein. Recently, it was shown that proteolytically truncated mouse JPH2 variants are imported into nucleus to exert alternative functions. However, the intranuclear behaviors of human JPH2 (hJPH2) and underlying molecular determinants have not been explored. Here, we demonstrate that full-length hJPH2 is imported into nucleus in human cells by two nuclear localization signals (NLSs), including a newly discovered one at the C-terminus. Importantly, unlike the JPH2 N-terminal truncation which diffuses throughout the nucleus, full-length hJPH2 forms nuclear bodies behaving like liquid-liquid phase separated droplets that are separated from chromatin. The C-terminal transmembrane domain is required for the formation of hJPH2 droplets. Oxidation mimicking substitution of residues C678 and M679 augments the formation of hJPH2 nuclear droplets, suggesting nuclear hJPH2 liquid-liquid phase separation could be modulated by oxidative stress. Mutation A405D, which introduces a negatively charged residue into an intrinsic disordered region (IDR) of hJPH2, turns liquid-like droplets into amyloid-like aggregates. Depletion of an Alanine Rich Region in the IDR recapitulates the liquid-amyloid phase transition. The MORN repeat regions of hJPH2 encodes intrinsic tendency to form amyloid-like structure. Together, these data for the first time revealed the novel intrinsic properties of hJPH2 to form nuclear liquid droplets, and identified critical functional domains and residues encoding these properties. We propose that hJPH2 droplets could function as membrane-less organelles participating in nuclear regulatory processes.

## Introduction

Junctophilin-2 (JPH2) is an essential protein whose loss of function results in life threatening heart diseases and early mortality of human [1–3]. Since the initial description of JPH2 as a membrane tethering protein bridging the plasma membrane (PM) and sarco/endoplasmic reticulum (SR/ER) in straited muscles [4], an extensive body of research has been focusing on its conventional function in stabilizing the junctional membrane complexes, where JPH2 binds membranes via its N-terminal Membrane Occupation and Recognition Nexus (MORN) repeats and a C-terminal transmembrane (TM) domain [4, 5]. Interestingly, recent studies with mouse JPH2 discovered that the JPH2 truncations can exert ‘moonlighting’ functions in nucleus under cardiac stresses that induce proteolytic cleavage at multiple sites on JPH2 [6, 7]. These studies converge in agreements that 1. Nuclear importation of JPH2 derivatives can result in functional transformation of this protein; 2. proteolytic cleavage that cuts off the C-terminal or N-terminal membrane binding domains is required to liberate the activity of a Nuclear Localization Signal (NLS) and trigger nuclear importation of JPH2 truncations. However, the sequence determinants of JPH2 nuclear importation revealed by those studies could be incomplete, because proteomic studies detected JPH2 in nuclear fraction under baseline condition [8, 9], suggesting it could be imported into nucleus by alternative mechanisms independent of stress induced proteolysis. Additionally, those discoveries were derived from mouse JPH2 [6, 7], which bears evolutionary divergency from human orthologue at several regions (Fig S1). The sequence determinants of human JPH2 (hJPH2) subcellular compartmentalization remain to be clarified.

Membrane-less organelles (MLOs) enclose specific proteins and nucleic acids to perform their functions within a confined space [10]. Liquid-liquid phase separation (LLPS) is the basis for the formation of MLOs in cells and is involved in many biological processes [10–12]. In many of these cases, phase separation is driven by multivalent interactions between intrinsically disordered proteins (IDPs) or their protein or nucleic acid partners, resulting in macromolecular condensates exhibit liquid-like properties [10–12]. There are multiple putative intrinsic disordered regions (IDRs) and modular hydrophobic regions throughout the hJPH2 protein (Fig S1), raising the question whether hJPH2 undergoes intracellular phase separation. The focus of this study is to determine the sequence determinants of hJPH2 subnuclear compartmentalization and phase separation.

## Material and methods

### Molecular cloning

cDNA of hJPH2 transcript (NM_020433.5) coding zone was synthesized by Lifesct.Inc, and cloned into pcDNA3.1 or a CMV promoter driven GFP fusion vector. A HA tag was fused to the C-terminus of hJPH2 cDNA. PCR based cloning and mutagenesis were used to construct the plasmids encoding the hJPH2 fusion proteins and mutations described in the context. All cloning products were verified by sanger sequencing (University of Minnesota genomic center). pLenti6-H2B-mCherry was a gift from Torsten Wittmann (Addgene plasmid # 89766) [13].

### Cell culture

Hela, 293T and HCT116 cells were purchased from ATCC and maintained in growth medium containing DMEM and 10% FBS. JetPRIME agent was used to transfect cells.

### Western blot and immunostaining

Western blotting and immunostaining were performed using antibody against hJPH2 (Santa Cruz, sc-398290) or HA tag (cell signaling, #3724). Nuclear fraction of Hela cells was harvested by ultracentrifugation of sucrose gradient as described previously [6]. Cells seeded on glass slips were fixed with 4% Paraformaldehyde and permeabilized with tritonX100. DAPI was used to stain nucleus. A Zeiss LSM900 was used to capture fluorescence images.

### Analysis of nuclear body images

The nuclear bodies captured in confocal images were automatically detected and analyzed by an in-house program developed using MATLAB (Fig. S2). The numbers, sizes and morphologies of nuclear bodies were quantified by the program.

### Live cell confocal imaging

Live cell time-lapse images were acquired on a Zeiss LSM900 confocal microscopy equipped with an environmental control system which maintained 37°C and 5% CO2 during the imaging. Transfected cells seeded on glass-bottomed chambers were maintained in growth medium during the imaging. For droplet fusion imaging, image sequences were acquired with an interval time of 1min. Fluorescence recovery after photobleaching (FRAP) analysis was performed in transfected 293T cells. Regions of interest were bleached using a laser intensity of 80% at 488nm. For diffusing proteins, or protein assemblies behaving like liquid droplets, recovery was recorded with an interval of 2~5s. For amyloid-like aggregates with slow molecule mobility, the intervals were set to be 5~10s. Images were processed by subtracting background. The flow of the protein was measured by quantifying the recovery of the bleached area at the cost of the unbleached region.

FRAP analysis was carried out using the following equation: F(t)=(I_bleach_(t)/I_total_(t))/(I_bleach_(t0)/I_total_(t0)), where I_bleach_ is the fluorescence intensity in the bleach area, Itotal is the fluorescence intensity of the entire condensate or nucleoplasm. 1-F(t) was fitted to 1^st^ exponential function to calculate time constant Ƭ. Interpolated time series data were averaged to show the overall behavior of specific GFP fused protein. The FRAP images were processed and analyzed by an in-house program developed using MATLAB.

### Sequence alignment and analysis

The alignment of mouse and human JPH2 orthologs was performed using Clustal Omega [14], and visualized using Jalview [15]. The putative IDRs were predicted by predictor VL-XT at www.pondr.com. The NLS^(619-657)^ was predicted by NLStradamus [16].

### Statistics

The histograms of droplet numbers per transfected nucleus were fitted to Poisson distribution. Kruskal-Wallis test or general linear model (GLM) were used for hypothesis tests when appropriate.

## Results

### Full-length hJPH2 forms nuclear bodies in human cells

Though JPH2 has been studied primarily in muscles, the Human Protein Atlas database shows that hJPH2 is also expressed in multiple human cell lines with various of origins [17]. The endogenous expression of hJPH2 protein in Hela cells (cervical cancer origin) is confirmed by western blot (Fig1A). Immunostaining using an antibody against a C-terminal epitope of hJPH2 showed that hJPH2 is distributed in both nuclear and extranuclear spaces throughout Hela cells (Fig 1B). In nucleus, hJPH2 forms spherical-like micron-scale bodies which are localized at foci with low DAPI staining intensity (Fig 1B), indicating they are confined at interchromatin spaces. To show whether these intranuclear assemblies of hJPH2 are specific to Hela cells, we expressed GFP fused hJPH2 containing a C-terminal HA tag (GFP-hJPH2-HA) in 293T cells. In both 293T cells and Hela cells, GFP-hJPH2-HA formed similar spherical intranuclear assemblies separated from chromatin DNA as indicated by DAPI staining (Fig 1C, Fig S3A), indicating it is an intrinsic property of hJPH2 to assemble into nuclear bodies. The extranuclear GFP-hJPH2-HA was localized to the PM and network-like structure representing ER. Coexpression of GFP-hJPH2-HA and histone H2B-mCherry, which labels the chromatin, confirmed that hJPH2 nuclear bodies are separated from chromatin (Fig S3B). Immunostaining against the C-terminal HA tag marked the same nuclear bodies as the N-terminal GFP did (Fig 1C), indicating those compartments are composed of full-length hJPH2 rather than truncations. This is consistent with the western blot data showing the presence of full-length hJPH2 rather than truncations in nuclear fraction of Hela cells (Fig 1A). To rule out the effect of GFP on hJPH2 condensation, we expressed hJPH2-HA in human cells, and immunostaining of HA tag showed similar nuclear bodies (Fig S3C&D).

**Fig1.**
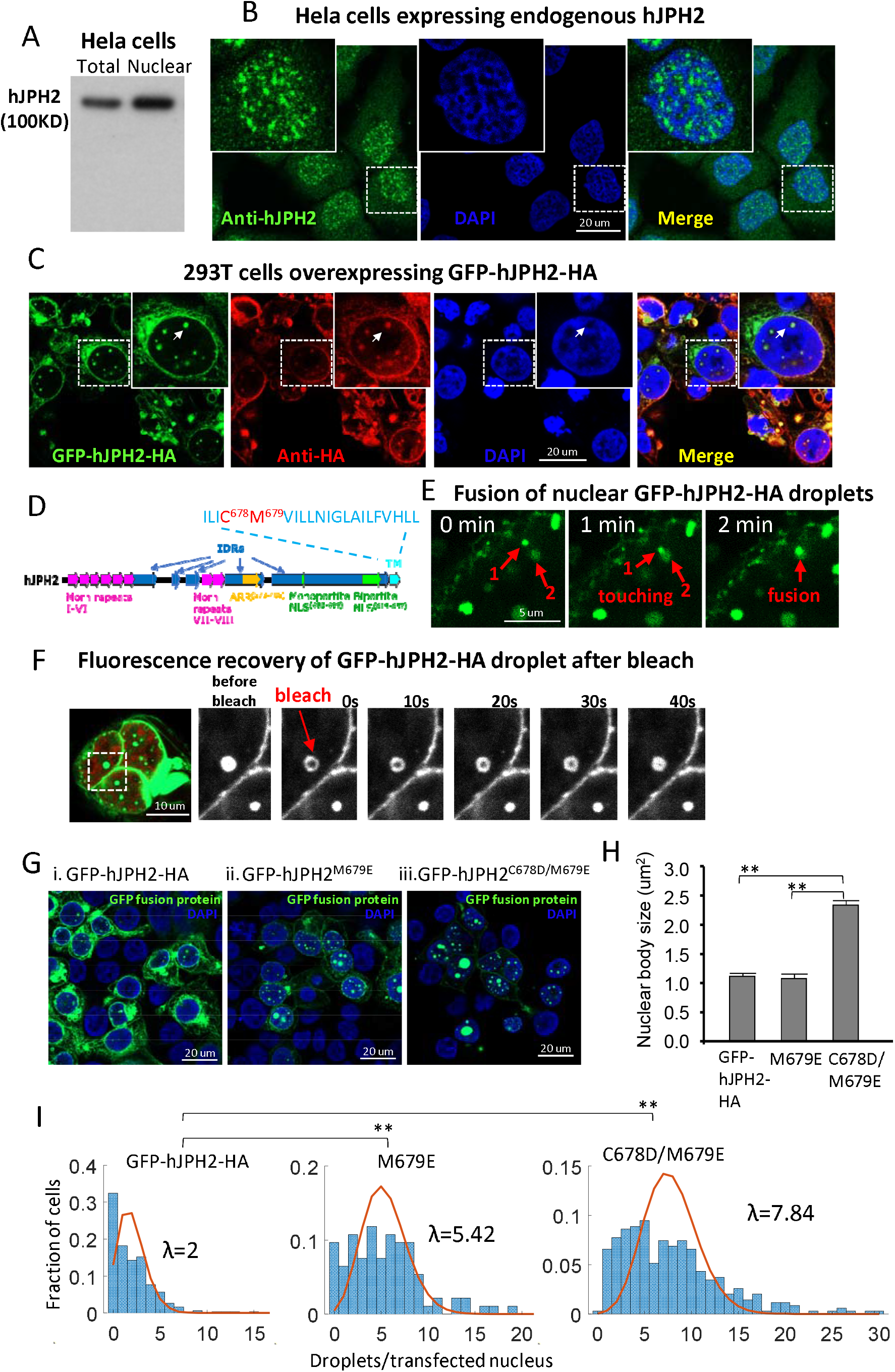
Full-length hJPH2 forms nuclear bodies that are phase separated liquids. A) Endogenous hJPH2 is expressed in Hela cells and distributed in nuclear fraction. B) Immunostaining showing endogenous hJPH2 nuclear bodies in Hela cells. C) GFP-hJPH2-HA forms nuclear bodies in 293T cells. White arrows indicate the separation of nuclear bodies from DNA (DAPI staining). D) Schematics of functional domains in hJPH2 protein. E) Time-lapse imaging showing fusion of two nuclear droplets (indicated by arrows) formed by GFP-hJPH2-HA in a Hela cell. F) FRAP of a nuclear droplet formed by GFP-hJPH2-HA in a 293T cell. Nuclear region is defined by co-transfected histone H2B-mCherry (red). G) Images of GFP-hJPH2-HA (i), mutant GFP-hJPH2^M679E^ (ii), and GFP-hJPH2^C678D/M679E^ (iii) in 293T cells. DAPI marks nucleus (blue). H) Sizes of nuclear droplets formed by GFP-hJPH2-HA, GFP-hJPH2^M679E^ and GFP-hJPH2^C678D/M679E^. ** p<0.01 by GLM. I) Histogram of nuclear droplet numbers in each transfected 293T cells. Data was fitted to Poisson distribution and λ (expected rate of occurrence) is calculated. ** p<0.01 by Kruskal-Wallis test. For H) and I), n= 612, 504, 2735 droplets from 302, 93, 349 cells, respectively.

### hJPH2 nuclear bodies behave as liquid droplets

Analysis of hJPH2 sequence revealed multiple putative IDRs, with the longest one at the C-terminal evolutionarily divergent region (Fig 1D, Fig S1). It is known that IDRs are excellent candidates for LLPS, which is responsible for the formation of intracellular liquid droplets that can function as MLOs. The morphology of nuclear hJPH2 assemblies is reminiscent of liquid droplets, which usually appear in spherical shape because of surface tension. Using time-lapse live cell imaging, we observed that intranuclear GFP-hJPH2-HA assemblies are mobile, and can fuse after touching one another (Fig.1E). Photobleaching analysis (FRAP) revealed that the GFP-hJPH2-HA molecules redistribute within the nuclear bodies with a time constant (Ƭ) of 114±15 s (Fig 1F, 2E&F). These behaviors agree with the typical hallmarks of liquid droplets [12]. Thus, hJPH2 nuclear bodies behave as liquid droplets.

It is known that a mouse JPH2 N-terminal truncation produced by proteolytic cleavage can be imported into nucleus because of the removal of nuclear exportation signal encoded by the TM domain [6]. However, the corresponding hJPH2 truncation hJPH2(1-572) (residue 1-572) is distinct from full-length hJPH2 by dispersing throughout the nucleus rather than forming droplets (Fig S4B). FRAP assay within nucleus expressing GFP-hJPH2(1-572) only detected minor local photobleaching, which recovered very rapidly (Fig 2D-vi, E, F), indicating that this truncated protein is highly diffusible and exists as a homogeneous solution in nucleoplasm. GFP-hJPH2(1-674), which lacks the TM domain, also dispersed in the nucleoplasm (Fig S4D). Therefore, the C-terminus TM domain of hJPH2 is required for the formation of liquid-droplets. Next, we searched the hJPH2 C-terminus for critical residues influencing the droplet formation.

### Mutations mimicking oxidized residues at C-terminus enhance formation of intranuclear hJPH2 droplets

Hydrophobic residues C678 and M679 are located at the beginning of the TM domain. GFP-hJPH2^M679E^, which bears a substitution of M679 for a hydrophilic residue, showed increased number of nuclear droplets (Fig 1G-ii, I). Mutation of both C678 and M679 into hydrophilic residues (C678D/M679E) further enhanced the numbers and sizes of nuclear droplets (Fig 1 G-iii, H, I). The liquid droplet characteristics such as inter-droplet fusion (Fig S5), intra-droplet molecule mobility (Fig 2D-i, E, F), and spherical morphology (Fig 2C) are preserved in C678D/M679E mutant. Cysteine and methionine residues are susceptible to oxidation. Given that the mutations C678D and M679E are oxidized residue mimetics, it is likely that oxidative stress may affect the intranuclear assembly of hJPH2 droplets.

**Fig2.**
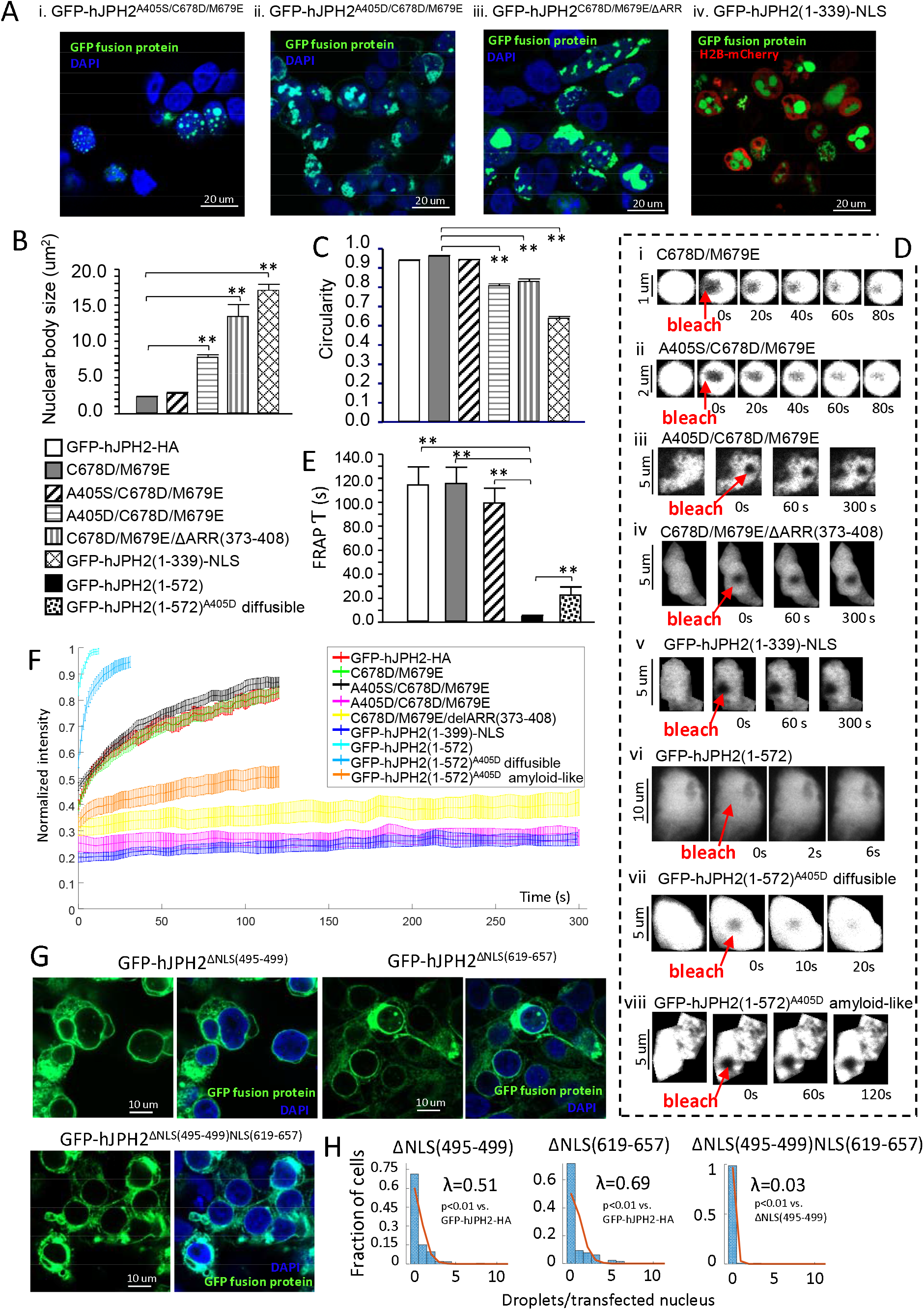
Sequence determinants of hJPH2 liquid-amyloid phase transition and nuclear localization. A) Nuclear bodies formed by GFP-hJPH2 mutants or truncations in 293T cells. Nucleus is marked by DAPI (blue) or H2B-mCherry (red). B-C) Sizes (B) and circularity (C) of nuclear bodies formed by indicated mutants in 293T cells. Circularity=1 represent a perfect round object. n=612, 2735, 807, 388, 156, 655 nuclear bodies from GFP-hJPH2-HA, GFP-hJPH2^C678D/M679E^, GFP-hJPH2^A405S/C678D/M679E^, GFP-hJPH2^A405D/C678D/M679E^, GFP-hJPH2^C678D/M679E/ΔARR(373-408)^, GFP-hJPH2(1-339)-NLS respectively. D) Examples of FRAP within nuclear bodies formed by GFP-hJPH2 mutants or truncations (i~v, viii), or in nucleoplasm dispersed with GFP-hJPH2(1-572) (vi) or diffusible GFP-hJPH2(1-572)^A405D^ (vii). Red arrowheads mark photobleaching regions. FRAP was performed in transfected 293T cells. E) Time constant (Ƭ) of molecule redistribution after photobleaching. F) Quantification of FRAP. Data are represented as Mean±SE for each time point. In E and F, n= 24, 41, 33, 11, 14, 10, 15, 40, 22 FRAP experiments for GFP-hJPH2-HA, GFP-hJPH2^C678D/M679E^, GFP-hJPH2^A405S/C678D/M679E^, GFP-hJPH2^A405D/C678D/M679E^, GFP-hJPH2^C678D/M679E/ΔARR(373-408)^, GFP-hJPH2(1-339)-NLS, GFP-hJPH2(1-572), GFP-hJPH2(1-572)^A405D^ diffusible, GFP-hJPH2(1-572)^A405D^ amyloid-like, respectively. G) Images of GFP-hJPH2 mutations lacking individual or both NLSs. H) Histogram of nuclear droplet numbers in each transfected 293T cells. n= 198, 127, 132 cells for GFP-hJPH2^ΔNLS(495-499)^, GFP-hJPH2^ΔNLS(619-657)^, GFP-hJPH2^ΔNLS(495-499)NLS(619-657)^, respectively. Significance between histograms were detected by Kruskal-Wallis test. ** p<0.01 by GLM in B, C and E.

### Mutations within an IDR of hJPH2 transformed nuclear liquid droplets into amyloid-like aggregates

The second longest IDR in hJPH2 overlaps with an Alanine Rich Region (ARR, residues 373-408) (Fig 1D, Fig S1). Depletion of the ARR from GFP-hJPH2_C678D/M679E_ resulted in formation of irregular shaped aggregates in nuclear and perinuclear regions (Fig 2A-iii). This phenotype can be recapitulated by introducing mutant A405D, which changes a nonpolar residue in the ARR into a negatively charged residue (Fig2A-ii). Compared with the GFP-hJPH2^C678D/M679E^ nuclear droplets, nuclear aggregates formed by GFP-hJPH2^A405D/C678D/M679E^ or GFP-hJPH2^C678D/M679E/ΔARR(373-408)^ are significantly larger and less spherical (Fig 2B, C). FRAP assay revealed low molecule mobility in these aggregates (Fig2D-iii&iv, E, F). These behaviors are consistent with amyloid-like state. Mutation A405S, which is an uncharged residue substitution, didn’t induce amyloid-like aggregation (Fig 2A-i, B~F).

A405D mutation also transformed rapidly diffusing GFP-hJPH2(1-572) into amyloid-like state in 22 out of 62 cells examined with FRAP (Fig2D-viii, E, F). In the rest cells, the diffusing rate of GFP-hJPH2(1-572)^A405D^ is much lower than that of GFP-hJPH2(1-572) (Fig 2D-vii, E, F).

hJPH2 MORN repeats (residues 1-339) fused with GFP and a SV40 NLS (GFP-hJPH2(1-339)-NLS) formed nuclear amyloid-like aggregates, as evidenced by large size, irregular shape and lacking of molecule mobility in the structure (Fig 2A-iv, B, C, D-v, F). These data show that the MORN repeat region of hJPH2 encodes intrinsic ability to form amyloid-like aggregates, and the ARR encodes a solubilizing domain neutralizing the amyloid tendency. Changing the charge of ARR can reduce its solubilizing effect.

### Identification of two NLSs that determine the nuclear localization of hJPH2

It is known that a monopartite NLS (KRPRP) is responsible for nuclear localization of mouse JPH2 truncations [6, 7]. Depletion of the corresponding NLS^(496-499)^ (residues 496-499) prevented the nuclear localization of GFP-hJPH2^(1-572)^ (Fig S4B&C). However, depletion of NLS^(496-499)^ didn’t prevent the nuclear localization of GFP-hJPH2^(1-674)^ (Fig S4D&E), indicating existence of redundant NLS at the C-terminus. By analysis of hJPH2 sequence, we identified a new bi-partite NLS^(619-657)^ (residues 619-657), which is localized at the C-terminal evolutionarily divergent region (Fig 1D, Fig S1). Depletion of both NLSs from GFP-hJPH2(1-674) prevented its nuclear localization (Fig S4F). Depletion of either of the two NLSs from GFP-hJPH2-HA significantly reduced the appearance of nuclear droplets (Fig 2G, H). Depletion of both NLSs from GFP-hJPH2-HA almost completely prevented the formation of nuclear droplets (Fig 2G, H).

## Discussion

This is the first study showing the liquid droplet like behaviors of full-length hJPH2 in nucleus. And this characteristic is not shared by the previously reported JPH2 N-terminal truncation, which diffuses quickly throughout the nucleoplasm. Many biological macromolecules, such as proteins and nucleic acids, exert their biological functions by forming phase-separated condensates, and phase separation is closely related to various human diseases [10, 12, 18]. We propose the hJPH2 droplets could function as MLOs involved in nuclear regulatory processes. These functions can be examined in future studies.

At subnuclear scale, hJPH2 droplets and chromatins appear to be mutually exclusive, though hJPH2 contains an ARR which was shown to have DNA binding activity [6]. However, that doesn’t rule out the possibility that hJPH2 droplet may establish stochastic interaction with loosely packed DNAs (euchromatic regions or interchromatin spaces). One such example is nuclear speckles, which are phase separated MLOs that are localized at transcription hot zones between chromatins [19]. Whether and how hJPH2 droplets affect gene expression can be examined in future studies.

This study for the first time shows the liquid-like to amyloid-like phase transition of full-length hJPH2 and JPH2 N-terminal truncation as a consequence of single residue mutation in an IDR. Detailed biophysical mechanisms underlying this property can be explored in future studies. A405 is identified as a critical residue determining the liquid phase of hJPH2. Interestingly, A405S is a human cardiac disease related mutation [3]. Though A405S alone doesn’t induce amyloid aggregation of hJPH2, the potential phosphorylation of Serine can mimic the A405D that induces liquid-amyloid transition. The detrimental effect of amyloid aggregates to cell viability may be a mechanism underlying the relationship between this mutation and disease.

Though JPH2 orthologs show high conservation in membrane tethering domains, they are evolutionarily divergent in IDRs which contain sequence determinants of nuclear localization and liquid property as shown in this study. Considering the critical roles of IDRs in LLPS, cautious interpretation is needed when translating JPH2 nuclear functions derived from mouse models to human physiology. To better define the functional consequences of nuclear hJPH2, genetic engineered mice with humanized JPH2 gene or human embryonic stem cell derived cell models can be used.

## 1. Abbreviations

JPH2: Junctophilin-2
hJPH2: human Junctophilin-2
LLPS: Liquid-Liquid Phase Separation
MLOs: Membrane-Less Organelles
FRAP: Fluorescence Recovery After Photobleaching
NLS: Nuclear Localization Signal
IDR: Intrinsic Disordered Region
IDP: Intrinsic Disordered Protein
ARR: Alanine Rich Region
MORN: Membrane Occupation and Recognition Nexus
TM: Transmembrane
PM: Plasma Membrane
SR/ER: Sarco/Endoplasmic Reticulum
GLM: General Linear Model

## Author Contributions

AG conceived and designed the study, developed the in-house programs for data analysis, analyzed the data, and wrote the manuscript. AG, WF and SG did the experiments.

## References

[1] E.G. Jones, N. Mazaheri, R. Maroofian, M. Zamani, T. Seifi, A. Sedaghat, G. Shariati, Y. Jamshidi, H.D. Allen, X.H.T. Wehrens, H. Galehdari, A.P. Landstrom, Analysis of enriched rare variants in JPH2-encoded junctophilin-2 among Greater Middle Eastern individuals reveals a novel homozygous variant associated with neonatal dilated cardiomyopathy, Sci Rep 9(1) (2019) 9038.

[2] S.U.M. Vanninen, K. Leivo, E.H. Seppälä, K. Aalto-Setälä, O. Pitkänen, P. Suursalmi, A.P. Annala, I. Anttila, T.P. Alastalo, S. Myllykangas, T.M. Heliö, J.W. Koskenvuo, Heterozygous junctophilin-2 (JPH2) p.(Thr161Lys) is a monogenic cause for HCM with heart failure, PLoS One 13(9) (2018) e0203422.

[3] A.P. Quick, A.P. Landstrom, Q. Wang, D.L. Beavers, J.O. Reynolds, G. Barreto-Torres, V. Tran, J. Showell, L.E. Philippen, S.A. Morris, D. Skapura, J.M. Bos, S.E. Pedersen, R.G. Pautler, M.J. Ackerman, X.H. Wehrens, Novel junctophilin-2 mutation A405S is associated with basal septal hypertrophy and diastolic dysfunction, JACC Basic Transl Sci 2(1) (2017) 56–67.

[4] H. Takeshima, S. Komazaki, M. Nishi, M. Iino, K. Kangawa, Junctophilins: a novel family of junctional membrane complex proteins, Mol Cell 6(1) (2000) 11–22.

[5] D.L. Beavers, A.P. Landstrom, D.Y. Chiang, X.H. Wehrens, Emerging roles of junctophilin-2 in the heart and implications for cardiac diseases, Cardiovasc Res 103(2) (2014) 198–205.

[6] A. Guo, Y. Wang, B. Chen, Y. Wang, J. Yuan, L. Zhang, D. Hall, J. Wu, Y. Shi, Q. Zhu, C. Chen, W.H. Thiel, X. Zhan, R.M. Weiss, F. Zhan, C.A. Musselman, M. Pufall, W. Zhu, K.F. Au, J. Hong, M.E. Anderson, C.E. Grueter, L.S. Song, E-C coupling structural protein junctophilin-2 encodes a stress-adaptive transcription regulator, Science (2018).

[7] S.K. Lahiri, A.P. Quick, B. Samson-Couterie, M. Hulsurkar, I. Elzenaar, R.J. van Oort, X.H.T. Wehrens, Nuclear localization of a novel calpain-2 mediated junctophilin-2 C-terminal cleavage peptide promotes cardiomyocyte remodeling, Basic Res Cardiol 115(4) (2020) 49.

[8] S. Franklin, M.J. Zhang, H. Chen, A.K. Paulsson, S.A. Mitchell-Jordan, Y. Li, P. Ping, T.M. Vondriska, Specialized compartments of cardiac nuclei exhibit distinct proteomic anatomy, Mol Cell Proteomics 10(1) (2011) M110 000703.

[9] A.A. Cutler, E.B. Dammer, D.M. Doung, N.T. Seyfried, A.H. Corbett, G.K. Pavlath, Biochemical isolation of myonuclei employed to define changes to the myonuclear proteome that occur with aging, Aging Cell 16(4) (2017) 738–749.

[10] S. Boeynaems, S. Alberti, N.L. Fawzi, T. Mittag, M. Polymenidou, F. Rousseau, J. Schymkowitz, J. Shorter, B. Wolozin, L. Van Den Bosch, P. Tompa, M. Fuxreiter, Protein Phase Separation: A New Phase in Cell Biology, Trends Cell Biol 28(6) (2018) 420–435.

[11] C.W. Pak, M. Kosno, A.S. Holehouse, S.B. Padrick, A. Mittal, R. Ali, A.A. Yunus, D.R. Liu, R.V. Pappu, M.K. Rosen, Sequence Determinants of Intracellular Phase Separation by Complex Coacervation of a Disordered Protein, Mol Cell 63(1) (2016) 72–85.

[12] D.M. Mitrea, R.W. Kriwacki, Phase separation in biology; functional organization of a higher order, Cell Communication and Signaling 14(1) (2016) 1.

[13] H. Pemble, P. Kumar, J. van Haren, T. Wittmann, GSK3-mediated CLASP2 phosphorylation modulates kinetochore dynamics, J Cell Sci 130(8) (2017) 1404–1412.

[14] F. Madeira, Y.M. Park, J. Lee, N. Buso, T. Gur, N. Madhusoodanan, P. Basutkar, A.R.N. Tivey, S.C. Potter, R.D. Finn, R. Lopez, The EMBL-EBI search and sequence analysis tools APIs in 2019, Nucleic Acids Res 47(W1) (2019) W636–W641.

[15] A.M. Waterhouse, J.B. Procter, D.M. Martin, M. Clamp, G.J. Barton, Jalview Version 2--a multiple sequence alignment editor and analysis workbench, Bioinformatics 25(9) (2009) 1189–91.

[16] A.N. Nguyen Ba, A. Pogoutse, N. Provart, A.M. Moses, NLStradamus: a simple Hidden Markov Model for nuclear localization signal prediction, BMC Bioinformatics 10 (2009) 202.

[17] M. Uhlén, L. Fagerberg, B.M. Hallström, C. Lindskog, P. Oksvold, A. Mardinoglu, Å. Sivertsson, C. Kampf, E. Sjöstedt, A. Asplund, I. Olsson, K. Edlund, E. Lundberg, S. Navani, C.A. Szigyarto, J. Odeberg, D. Djureinovic, J.O. Takanen, S. Hober, T. Alm, P.H. Edqvist, H. Berling, H. Tegel, J. Mulder, J. Rockberg, P. Nilsson, J.M. Schwenk, M. Hamsten, K. von Feilitzen, M. Forsberg, L. Persson, F. Johansson, M. Zwahlen, G. von Heijne, J. Nielsen, F. Pontén, Proteomics. Tissue-based map of the human proteome, Science 347(6220) (2015) 1260419.

[18] S. Alberti, D. Dormann, Liquid-Liquid Phase Separation in Disease, Annu Rev Genet 53 (2019) 171–194.

[19] Y. Chen, Y. Zhang, Y. Wang, L. Zhang, E.K. Brinkman, S.A. Adam, R. Goldman, B. van Steensel, J. Ma, A.S. Belmont, Mapping 3D genome organization relative to nuclear compartments using TSA-Seq as a cytological ruler, J Cell Biol 217(11) (2018) 4025–4048.

