## supplemental Figures for "Sequence Determinants of Human Junctophilin-2 Protein Nuclear Localization and Phase Separation"

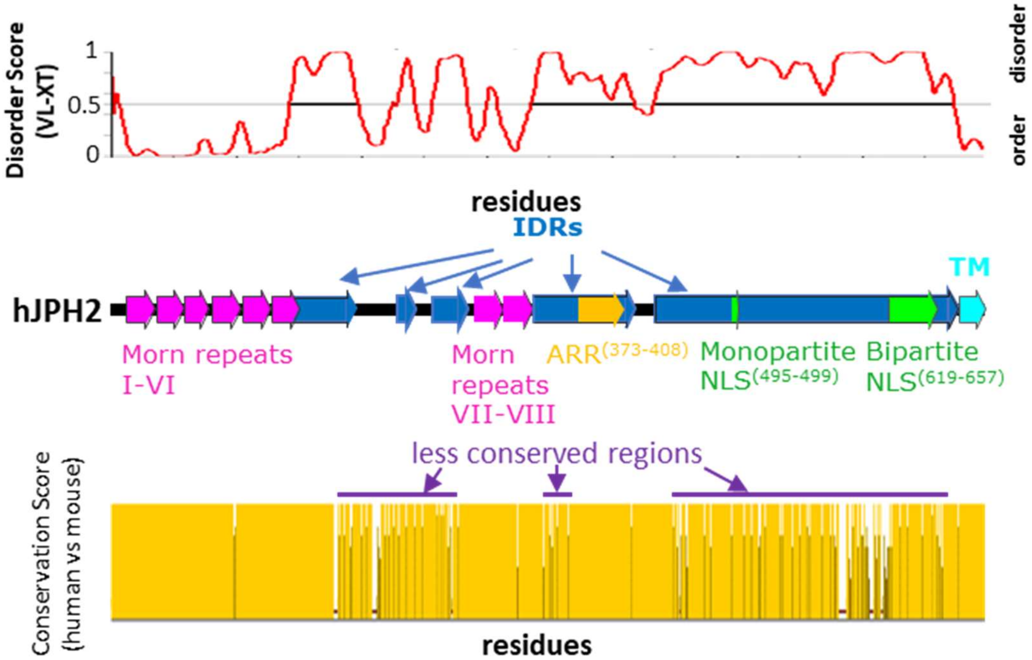


Fig S1. Schematic representation of intrinsic disordered regions (IDRs) and evolutionarily less conserved regions in hJPH2 protein.

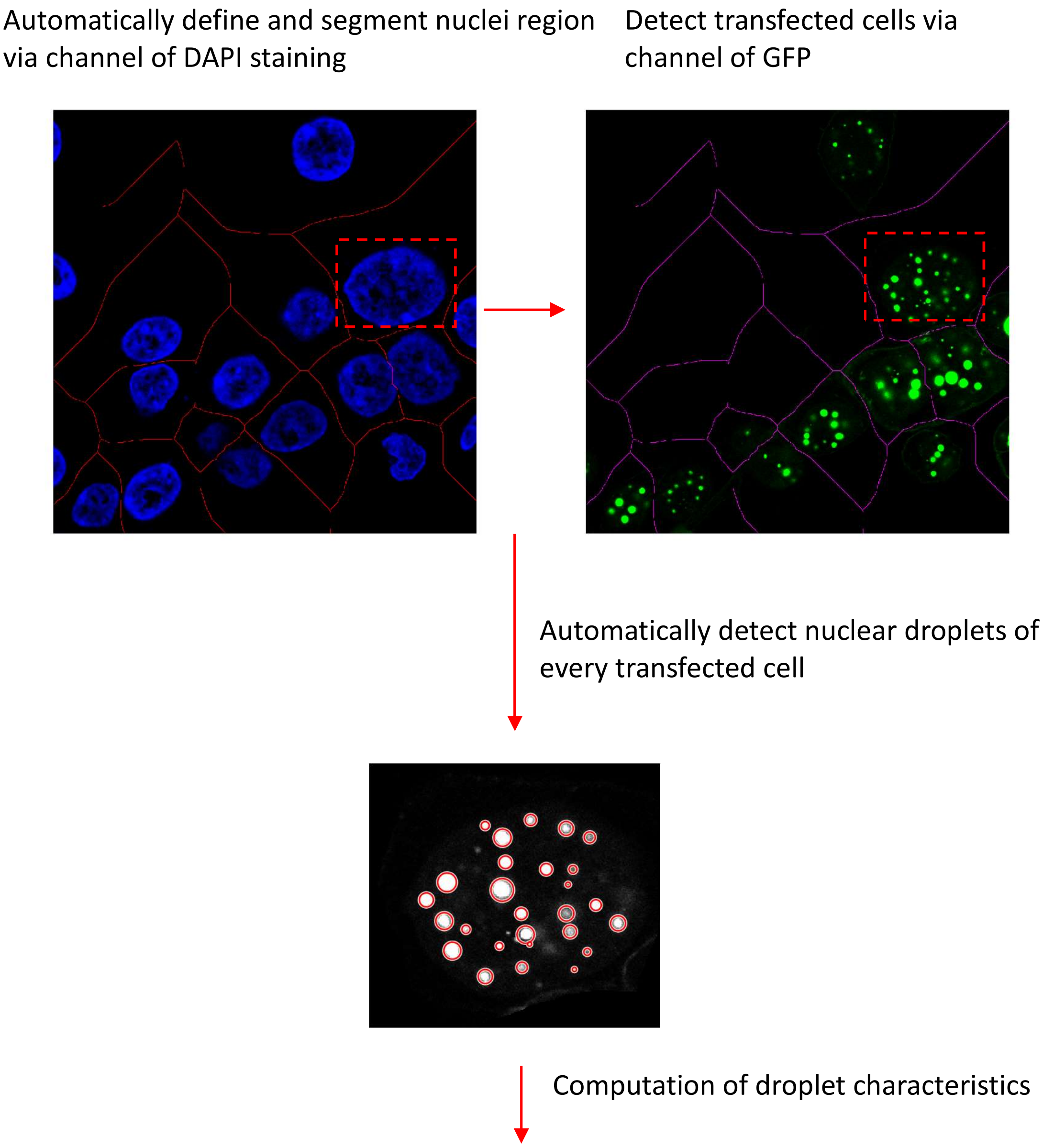


| | A | B | C | D | E | F | G | H | I | J | K | L | M | N | O |
|---|---|---|---|---|---|---|---|---|---|---|---|---|---|---|---|
| 1 | Area | Centroid_1 | Centroid_2 | MajorAxisLength | MinorAxisLength | Circularity | WeightedCentroid_1 | WeightedCentroid_2 | MeanIntensity | MinIntensity | MaxIntensity | Integratedsignal | Diameter | Number | CellName |
| 2 | 182 | 62.37912088 | 142.0549451 | 16.20878282 | 14.48343277 | 1.030254276 | 62.39683589 | 141.9956789 | 232.6923077 | 129 | 255 | 42350 | 15.34610779 | 1 | cell1 |
| 3 | 327 | 81.26299694 | 120.4495413 | 21.28367778 | 19.64419359 | 1.05929902 | 81.21450699 | 120.4624653 | 249.7584098 | 164 | 255 | 81671 | 20.46393569 | 2 | cell1 |
| 4 | 21 | 74.38095238 | 177.5238095 | 5.697594671 | 5.020592515 | 1.249793188 | 74.64767932 | 177.6181435 | 22.57142857 | 12 | 40 | 474 | 5.359093593 | 3 | cell1 |
| 5 | 5 | 85.4 | 88.8 | 3.306559138 | 2.12916259 | 2.551915845 | 85.32142857 | 88.82142857 | 22.4 | 12 | 34 | 112 | 2.717860864 | 4 | cell1 |
| 6 | 305 | 94.88852459 | 161.8163934 | 20.23348189 | 19.32458499 | 1.033619508 | 94.85813117 | 161.7922311 | 249.757377 | 153 | 255 | 76176 | 19.77903344 | 5 | cell1 |
| 7 | 322 | 112.3571429 | 138.0279503 | 21.396071 | 19.2754246 | 1.043369803 | 112.3612038 | 138.0261575 | 248.3757764 | 155 | 255 | 79977 | 20.3357478 | 6 | cell1 |
| 8 | 93 | 107.7419355 | 82.92473118 | 12.01380495 | 10.03094285 | 1.098444594 | 107.6732525 | 82.98779678 | 129.5268817 | 41 | 255 | 12046 | 11.0223739 | 7 | cell1 |
| 9 | 44 | 128.1363636 | 171.2272727 | 8.958445468 | 6.481992537 | 1.191269053 | 128.0794045 | 171.2692308 | 18.31818182 | 0 | 41 | 806 | 7.720219002 | 8 | cell1 |
| 10 | 190 | 135.9052632 | 93.01578947 | 16.20446111 | 15.0360196 | 1.106412908 | 135.8969396 | 92.99696074 | 249.3684211 | 115 | 255 | 47380 | 15.62024035 | 9 | cell1 |
| 11 | 95 | 137.9157895 | 155.4842105 | 25.52353501 | 7.882255699 | 0.366601153 | 137.9360165 | 154.9035088 | 20.4 | 0 | 47 | 1938 | 16.70289535 | 10 | cell1 |
| 12 | 160 | 147.34375 | 125.725 | 15.45951618 | 13.39661902 | 1.032807698 | 147.1374831 | 125.5345305 | 88.05625 | 34 | 162 | 14089 | 14.4280676 | 11 | cell1 |
| 13 | 124 | 93.12903226 | 149.6693548 | 14.50320088 | 11.15225262 | 0.996685682 | 93.4609972 | 149.0376787 | 106.5887097 | 21 | 255 | 13217 | 12.82772675 | 1 | cell2 |
| 14 | 161 | 96.2173913 | 127.8757764 | 15.6190759 | 13.45962493 | 0.858123775 | 96.16432755 | 127.7196867 | 179.2360248 | 62 | 255 | 28857 | 14.53935041 | 2 | cell2 |
| 15 | 33 | 96.60606061 | 90.48484848 | 6.78219459 | 6.334130731 | 1.352853371 | 96.73186813 | 90.32307692 | 13.78787879 | 0 | 36 | 455 | 6.558162661 | 3 | cell2 |

Fig S2. Algorithm of the in-house program that automates nuclear droplet detection and quantification.

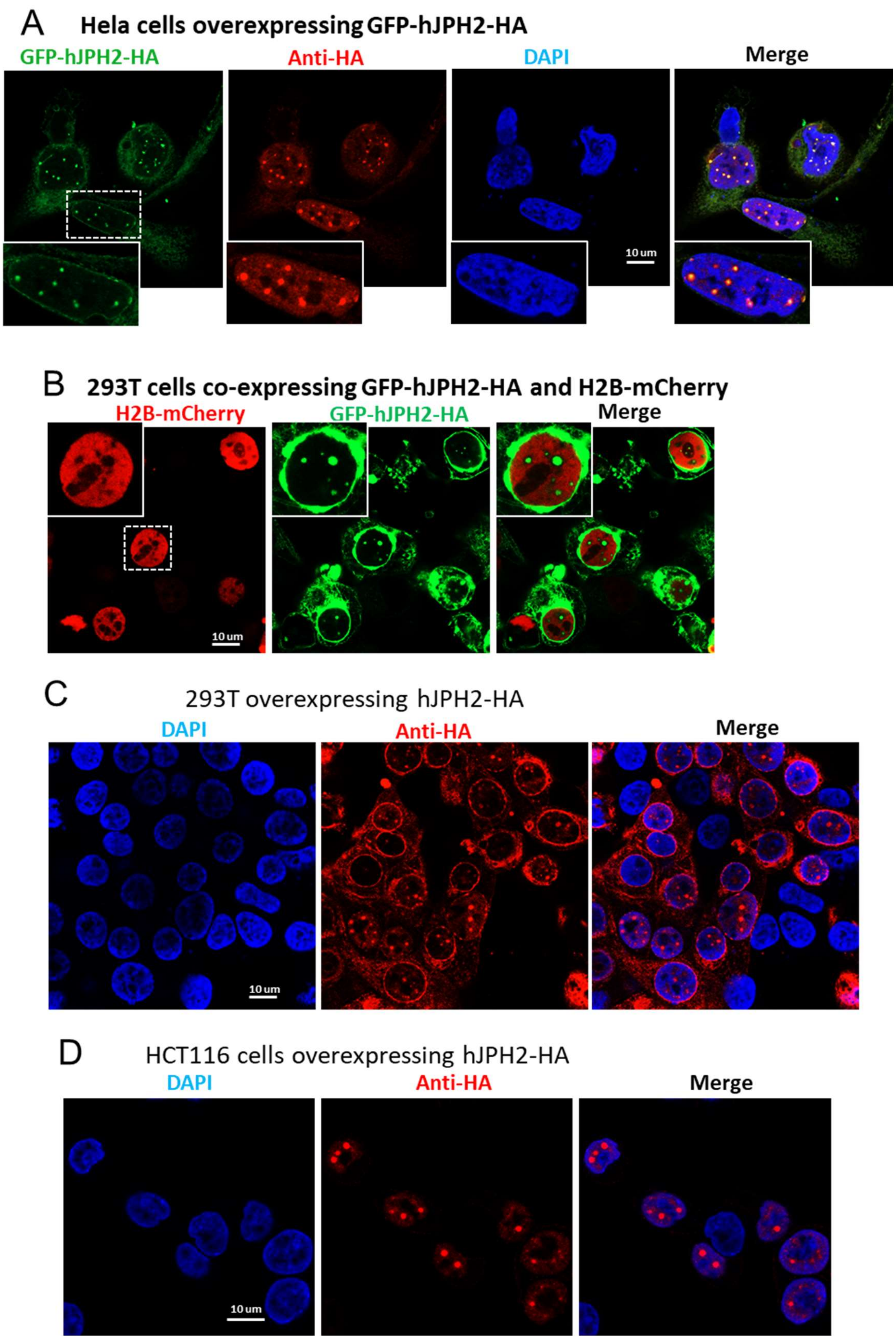


Fig S3. hJPH2 forms nuclear droplets in human cells. A) Immunostaining images of Hela cells overexpressing GFP-hJPH2-HA. B) Images of 293T cells co-expressing GFP-hJPH2-HA and histone H2B-mCherry, which labels the chromatin. Note that GFP-hJPH2-HA droplets are separated from chromatin. C) Immunostaining images of 293T cells overexpressing hJPH2-HA. D) Immunostaining images of HCT116 cells overexpressing hJPH2-HA.

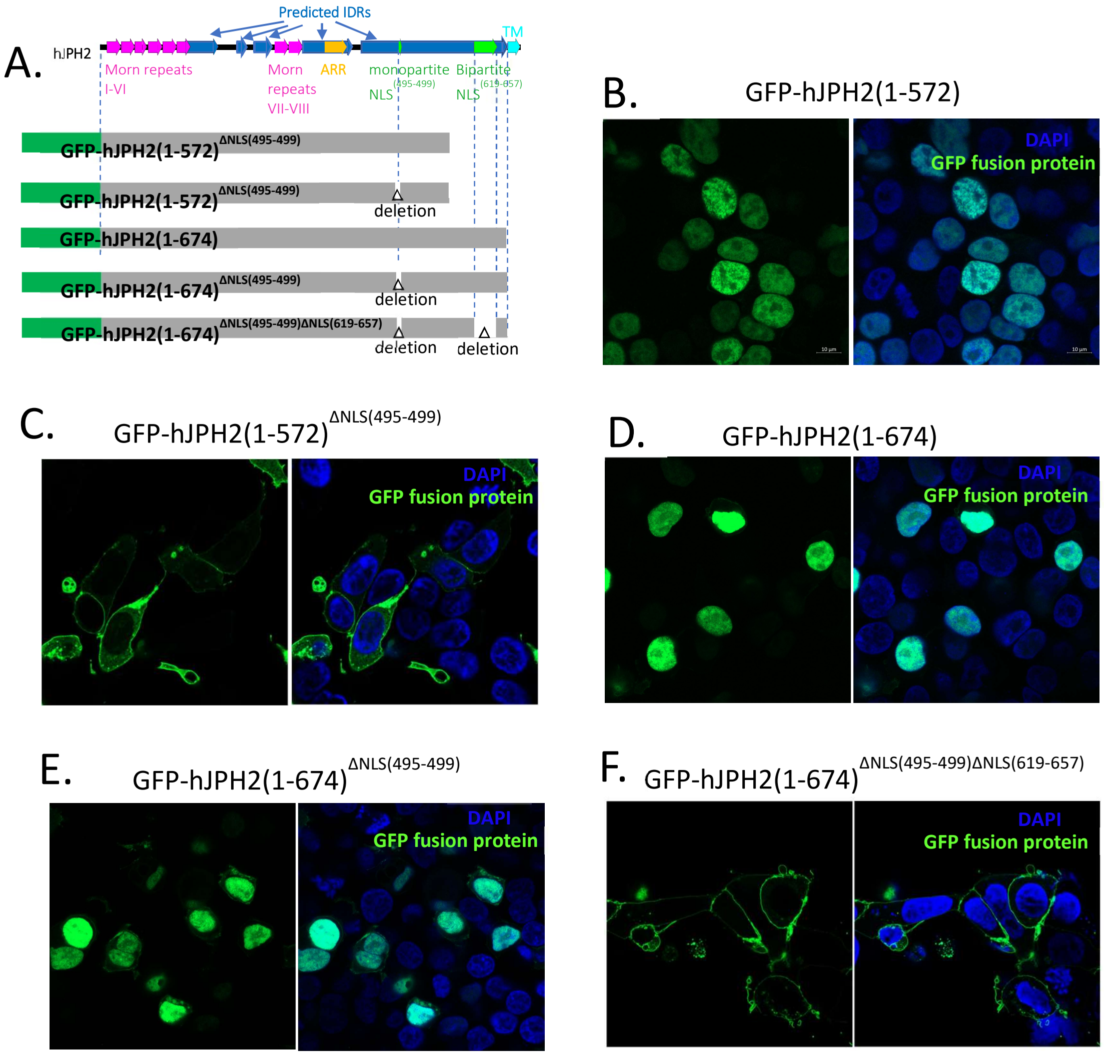


Fig S4. hJPH2 contains two NLSs. A) Schematics of fusion proteins used in these experiments. B~F) Images of 293T cells expressing GFP-hJPH2(1-572) and GFP-hJPH2(1-674) truncations with depletion of NLSs as indicated. Nucleus is marked by DAPI in blue. Note that depletion of NLS$^{(495-499)}$ doesn't prevent nuclear localization of GFP-hJPH2(1-674). Depletion of both NLSs prevented the nuclear localization of that protein.

**Fusion of GFP-hJPH2$^{C678D/M679E}$ nuclear droplets in a Hela cell**

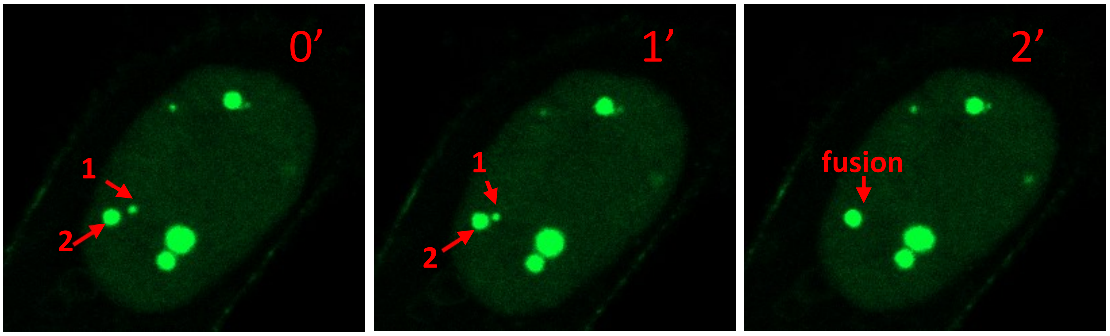


Fig S5. Time lapse imaging of fusion of nuclear droplets formed by GFP-hJPH2$^{C678D/M679E}$ in a Hela cell.
